# Latent *Toxoplasma gondii* infection increases soluble mutant huntingtin and promotes neurodegeneration in the YAC128 mouse model of Huntington’s disease

**DOI:** 10.1101/550624

**Authors:** David W. Donley, Teal Jenkins, Cailin Deiter, Reed Campbell, Marley Realing, Vanita Chopra, Stephen Hersch, Jason P. Gigley, Jonathan H. Fox

## Abstract

*Toxoplasma gondii* causes a prevalent neuroinvasive protozoal pathogen that in immune competent individuals results in latent infection characterized by intra-cellular parasite cysts in brain. Despite life-long infection, the role of latent toxoplasmosis on chronic neurodegenerative processes is poorly understood. Huntington’s disease (HD) is a progressive neurodegenerative disorder caused by a dominant CAG repeat expansion in the huntingtin gene (HTT) that results in the expression and accumulation of mutant huntingtin protein (mHTT). The mutant HD gene is fully penetrant. However, there is significant variability in disease progression that is in part explained by as yet unidentified environmental factors. The kynurenine pathway of tryptophan metabolism (KP) is an inflammatory pathway and its activation is implicated in HD pathogenesis. KP upregulation also occurs in response to infection with *Toxoplasma gondii* suggesting that the latent infection may promote HD. We discovered that mice on the FVB/NJ background develop latent toxoplasmosis following infection with the ME49 strain of *T. gondii*. This finding enabled us to address the hypothesis that latent toxoplasmosis potentiates disease in the YAC128 mouse model of HD, as these mice are maintained on the FVB/NJ background. Wild-type and HD mice were infected at 2-months of age. During the 10-month follow-up, infection had adverse effects on mice of both genotypes. However, YAC128 HD mice demonstrated specific vulnerability to latent toxoplasmosis, as demonstrated by the presence of increased striatal degeneration, high levels of the blood neurodegeneration marker neurofilament light protein, and elevated brain soluble mHTT. Our studies have uncovered a novel HD-infection interaction in mice that provides insights into the large variability of the human HD phenotype.

## Introduction

Huntington’s disease (HD) is a progressive autosomal dominant neurodegenerative disorder caused by a polyglutamine-encoding CAG-repeat expansion in the huntingtin gene (HTT) [1]. Mutant huntingtin protein (mHTT) accumulates in neurons and glial cells, resulting in transcriptional dysregulation, altered mitochondrial function, inflammation, and oxidative stress that promote neurodegenerative processes in the HD brain [2–4]. As mHTT is the upstream driver of HD, decreasing the level of the mutant protein is protective, and increasing mHTT levels is predicted to promote disease progression. Human HD is characterized by high variability in the age of onset after adjustment for CAG repeat length size [5]. Environmental factors account for ~60% of the non-CAG variability in the age of onset, with the remaining effect explained by non-HTT genetic factors [5]. Currently, environmental factors that modulate HD progression are unknown.

*Toxoplasma gondii* is a protozoan parasite with a human prevalence of ~20-30% in most developed countries, but infection rates are up to 80% in some countries [6–8]. The acute infection is characterized by proliferation of the tachyzoite stage of the parasite and the primary immune response. Even with an effective immune response, the parasite evades complete clearance, forming a latent infection characterized by intra-cellular cysts in the brain and striated muscles [9–11]. Seropositivity for *T. gondii* is associated with several neuropsychiatric disorders including schizophrenia, depression, and generalized anxiety [12–16]. However, meta-analyses and broader cohort studies report conflicting results, with some studies reporting strong correlation with neuropsychiatric disease and *T. gondii* infection in humans, and others finding no association [16–21]. Similarly, in mice, *T. gondii* infection exacerbates beta-amyloid (Aβ)-induced memory deficits but is associated with a reduction in Aβ load and decreased neuronal degeneration in the hippocampus [22–25].

HD patients demonstrate immune activation beginning early in the disease. Pre-clinically there is microglial activation in brain and the blood cytokines IL-6, IL-8 and TNFα are elevated [26, 27]. Mutant HTT levels in monocytes and T lymphocytes correlate with disease burden and striatal atrophy in human HD patients [28]. Monocytic mutant *HTT* impairs cell migration in response to chemotactic stimuli [24]. However, whether the inflammatory processes in HD are intrinsic responses to mHTT or represent altered responses to immune challenge by infection are not known. The kynurenine pathway of tryptophan metabolism (KP) is activated by inflammatory stimuli including IFNγ and IL-6 [29, 30]. This pathway is implicated in HD progression in mouse models [31–33]. Additionally, KP inhibition provides protection in HD models [34, 35]. *T. gondii* is a tryptophan auxotroph, so utilizes host tryptophan for growth. KP activation provides protection against the infection because it results in tryptophan depletion [30, 36, 37]. Therefore, *T. gondii* infection activates a pathway that is linked with HD potentiation.

Whereas the role of aseptic inflammation in HD and related protein-misfolding diseases has been well studied [26–28], how common latent brain infections, such as with *T. gondii* or HSV1, affect this and the underlying diseases is poorly understood. Interestingly, a transgenic mouse model of Alzheimer’s disease has an enhanced inflammatory response to acute *T. gondii* infection [38]. We have previously studied *T. gondii* infection in N171-82Q HD mice and reported premature mortality compared to non-infected HD mice [39]. However, on the B6/C3H genetic background of these mice, latent infection was not established, and the mice died before effects on HD could be investigated. We have since discovered that FVB/NJ background mice have high-genetic resistance to infection and establish long-term latent infection, thus enabling the current investigation in YAC128 HD mice. We now report that latent *T. gondii* infection in YAC128 HD mice promotes striatal degeneration, increases markers of KP activity, and increases brain mHTT levels. The significance of these results is that, in contrast to being designated as a silent infection, *T. gondii* increases established proximal and distal markers of HD in a mouse model. We also show that latent toxoplasmosis also promotes cortical degeneration in wild-type mice. These findings are relevant to understanding a potential link between latent toxoplasmosis and neurodegenerative processes in HD and aging.

## Materials and Methods

### Mouse husbandry and breeding

The YAC128 model is a transgenic model of HD that expresses full-length human mHTT. These mice display a progressive phenotype with mild disease, including behavioral deficits and neurodegeneration present before 1-year of age. They develop striatal and neocortical degeneration, as occurs in human HD [40]. WT and YAC128 HD mice (Jackson Labs) were maintained as described [41]. Mice were housed in OptiMice cages (Animal Care Systems) with 4-5 per cage and one genotype/treatment per cage [39]. Experimental mice were obtained by breeding mutant *HTT* males with WT females. Sentinel mice were evaluated every 6 months by comprehensive serology panel screening for murine infectious diseases at Charles River Laboratories and were free from all tested diseases. All mice were checked daily. They were sacrificed using B-euthanasia solution.

### Ethics Statement

This study and all procedures were carried out in accordance with the National Institutes of Health guidelines for animal welfare and were approved by the University of Wyoming Institutional Animal Care and Use Committee (UW-IACUC) (protocol numbers 20150929JG00198-03 and 20180129JF00292-01).

### Experimental Design

Male and female YAC128 HD mice were used for all experiments. HD and WT littermate mice were weaned at 3.5 weeks of age and systematically assigned to experimental groups to balance ages and minimize effects of litter of origin. HD and WT mice were dosed with 10 Me49 strain *T. gondii* parasites cysts by oral gavage at 2-months of age. Experiments were a 2 × 2 factorial design. For all studies, the investigators were blinded with respect to the treatment and genotype, until data analysis. Individual mice were identified by a 3-digit identifier assigned during genotyping. Experiments were terminated at ~1 year of age, at which all mice were humanely sacrificed. All mice were accounted for in all studies. Mouse deaths prior to sacrifice are included in (**S1 Table**).

### *T. gondii* infections

The Me49 strain of *T. gondii* was harvested from young FVB/NJ mice infected with 10 cysts by oral gavage and sacrificed 4-5 weeks post-infection. Brain cysts were collected, counted, and suspended in cold PBS at a concentration of 50 cysts/ml. Experimental mice were infected once with 10 parasite cysts (200 μl) by oral gavage. Control mice received PBS by oral gavage.

### Mouse rotarod

Performance on the rotarod was completed as described [41].

### Neurofilament light chain ELISA

Neurofilament light chain (NFL) was measured in plasma at 8 and 12 months of age. Blood was collected from cardiac puncture and placed in a tube with 5 μl of 1 mM EDTA. Plasma was stored in liquid nitrogen until analysis. NFL ELISA was performed according to the manufacturer’s instructions (MyBioSource). Undiluted plasma (50 μl) was added to each well in duplicate and incubated for 2 hours. NFL concentrations were calculated using standards.

### Kynurenine pathway metabolite analyses

Plasma was collected from the retro-orbital sinus. In brief, mice were lightly anesthetized with an intraperitoneal injection of 30 mg/kg ketamine and 3.3 mg/kg xylazine; one drop of proparacain was used as a topical anesthetic at the collection site just before the procedure. After collection, mice were placed back in the home cage and closely monitored until fully recovered. Collected plasma was frozen in liquid nitrogen until analysis. Samples were diluted 1:3 (v/v) with methanol containing 2% (v/v) acetic acid and centrifuged at 12000 ×g for 5 minutes at 4°C. Supernatants were filtered using a Phenomenex Phree Phospholipid extraction column and a 0.2-μm filter. The samples were then run on a Waters Acquity UPLC-MS/MS with a methanol/2% acetic acid mobile phase gradient on a 2 × 100-mm Waters BEH C^18^ column. Each sample (5 μl) was injected with a flow rate of 0.3 ml/minute and a total run time of 4.5 minutes. Analytes were verified using two M+H parent-daughter transitions: tryptophan parent m/z = 205.13 to daughter m/z = 118.03 and 146.07, kynurenic acid parent m/z = 190.07 to daughter m/z = 88.96 and 116.03, kynurenine parent m/z = 209.04 to daughter m/z = 94.02and146.03, and 3-HK parent m/z = 224.98 to daughter m/z = 110.04 and 162.05. Concentrations were quantified using QuanLynx (Waters) software from serial dilutions of standards.

### Quantitative neuropathology

Anaesthetized mice were perfused with 4% (w/v) paraformaldehyde as described [17, 41]. Fixed brains were sectioned at 40 μm using a freezing microtome, and serial sections were collected in 12-well plates. Every 6^th^ section was mounted and stained with thionin. The Cavalieri and optical fractionator methods in StereoInvestigator (MicroBrightField) software were used to determine brain region volume and neuronal estimates, respectively. The optical fractionator method was applied to the striatum as described [42], whereas application to the neocortex used a 550 × 550-μm grid size and 27 × 27 × 10-μm counting frame. Cortical and striatal evaluations were completed on every twelfth serial section, and hippocampal evaluation was completed on every sixth serial section. Striatal and hippocampal analyses were performed on the entire structure, whereas the cerebral neocortex was quantified from the most rostral aspect of the lateral ventricle and terminated at the rostral hippocampus.

### Functional assessment of CD8+ T lymphocytes *ex vivo*

Assessment of antigen-specific IFNγ production from CD8+ T cells was performed as described [39]. Flow cytometry data were collected using Guava easyCyte HT flow cytometer (Millipore). Data were analyzed using FlowJo v9.3 software. Cell gating and the compensation strategy were determined from no-stain, single-color, and fluorescence-minus-one controls for each antibody.

### Quantitative real-time PCR analyses

Total RNA was extracted from cerebral cortex and striatum using phenol/chloroform and then purified using a column-based method (RNAeasy, Qiagen) with on-column DNase I digestion. *mHTT* was amplified using forward (5’-GTGCTGAGCGGCGCCGCGAGTC-3’) and reverse (5’-GGACTTGAGGGACTCGAAGGC-3’) primers that specifically target the mutant gene according to a previously reported protocol [43]. Expression was quantified using SYBR Green Master Mix (Life Technologies) and was normalized to β-actin using the Applied Biosystems Taqman gene expression primer/probe combination Mm00607939_s1. Expression analyses of *SAG1, BAG1*, and the *T. gondii* actin gene were performed as described using SYBR Green Master Mix and the Pfaffl method [44]. *SAG1* and *BAG1* expression is reported as a ratio relative to parasite actin. Gene expression was determined using 20 ng of cDNA per reaction for *mHTT* and 40 ng of cDNA per reaction for *SAG1* and *BAG1*. Amplifications used a Bio-Rad CFX real-time thermal cycler. *T. gondii* burden was determined by quantitative PCR on genomic DNA using DNA standards as described [39]. Reactions were carried out with 400 ng of genomic DNA purified brain homogenate using a Qiagen DNeasy Kit.

### Soluble mutant huntingtin

Soluble mHTT was measured by time-resolved fluorescence resonance energy transfer (TR-FRET) using antibodies targeted specifically to the N-terminal region of the mutant protein. Measurements were completed as described [45].

### Statistical analyses

All data were analyzed using SAS software version 9.2 (Cary, NC). Generalized linear modeling was used for single-time point analyses, and the mixed procedure was used for repeated-measures analyses. Assumptions of normality and equal variance were verified for each experiment. We investigated all main effects and interactions in the initial statistical model, and then performed pre-planned pairwise comparisons. Data that were not normally distributed were log transformed before analysis. These transformed data are presented as the mean ± 95%confidence interval (CI). P < 0.05 were considered significant.

## Results

### Latent *T. gondii* exacerbates HD-induced neurodegeneration

Preliminary evaluation of ME49 strain *T. gondii* infection at 2 months in FVB/J mice resulted in the fortuitous finding that the mice established a persistent latent (sub-clinical) infection without overt disease or death up to 12-months of age. YAC128 HD mice are maintained on this line enabling us to evaluate how *T. gondii* infection impacts HD in this mouse model. Subsequent experiments described here, where mice were infected orally with 10 *T. gondii* cysts at 2 months, corroborate these findings as we did not detect any differences in mortality due to infection or genotype up to 12-months of age (**Table S1, Fig. S1**).

Neurofilament light chain (NFL) is a blood marker of active neurodegeneration in HD [46, 47] and is also elevated in other neurodegenerative disorders [48]. We quantified blood NFL at 8 and 12 months of age to determine the effects of both infection and HD. NFL levels were increased by *T. gondii* infection (F_(1,25)_ = 35.5, p < 0.0001) and HD (F_(1,25)_ = 12.83, p = 0.0014) at 8 months. Similarly, at 12 months NFL was also increased by both *T. gondii* infection (F_(1,50)_ = 18.7, p < 0.0001) and by HD (F_(1,50)_ = 10.2, p = 0.0024). Infected HD mice had significantly increased NFL compared with both infected wild-type and non-infected HD mice (**Fig. 1A**). We then completed comprehensive quantitative neuropathology studies in 12-month-old mice to assess neurodegeneration directly (**Fig. 1B-H**). Brain weights were significantly decreased by infection (F_(1,54)_ = 71.25, p < 0.0001) and HD (F_(1,54)_ = 12.15, p = 0.001); there was also a significant genotype-infection interaction (F_(1,54)_ = 4.8, p = 0.0329) (**Fig. 1C**). The striatum preferentially degenerates early in HD models and human HD. Therefore, we quantified striatal volumes and estimated the number of striatal neurons (Figs. 1D-E). *T. gondii* infection (F_(1,40)_ = 4.96, p = 0.0316) and HD (F_(1,40)_ = 30.64, p < 0.0001) both significantly decreased striatal volumes (Figs. 1B, D). We also found a significant HD-T. *gondii* interaction (F_(1,40)_ = 6.2, p = 0.017), demonstrating that infected HD mice had significantly smaller striata compared with infected wild-type and non-infected HD mice (**Fig. 1D**). Consistent with this, unilateral total striatal neuronal estimates were decreased by *T. gondii* (F_(1,40)_ = 4.23, p = 0.0462) and HD (F_(1,40)_ = 5.37, p = 0.0257). Infected, versus non-infected, HD mice had significantly fewer striatal neurons (Fig 1E). Cerebral cortex undergoes extensive degeneration in HD models and human HD. We determined both neocortical volume and neuronal estimates (Figs. 1F-G). Interestingly, *T. gondii* promoted neocortical atrophy (F_(1,39)_ = 7.21, p = 0.0106) in WT and HD mice combined. Pair-wise comparisons were not significant however (**Fig. 1F**). Consistent with this, *T. gondii*infection resulted in loss of neocortical neurons (F_(1,39)_ = 13.46, p = 0.0007) in both WT and HD mice (**Fig. 1G**). To address the specificity of the infection to HD vulnerable brain areas we assessed hippocampal volumes as the hippocampus does not degenerate early in HD. There were no significant differences between the four groups (**Fig. 1H**).

**Figure 1.**
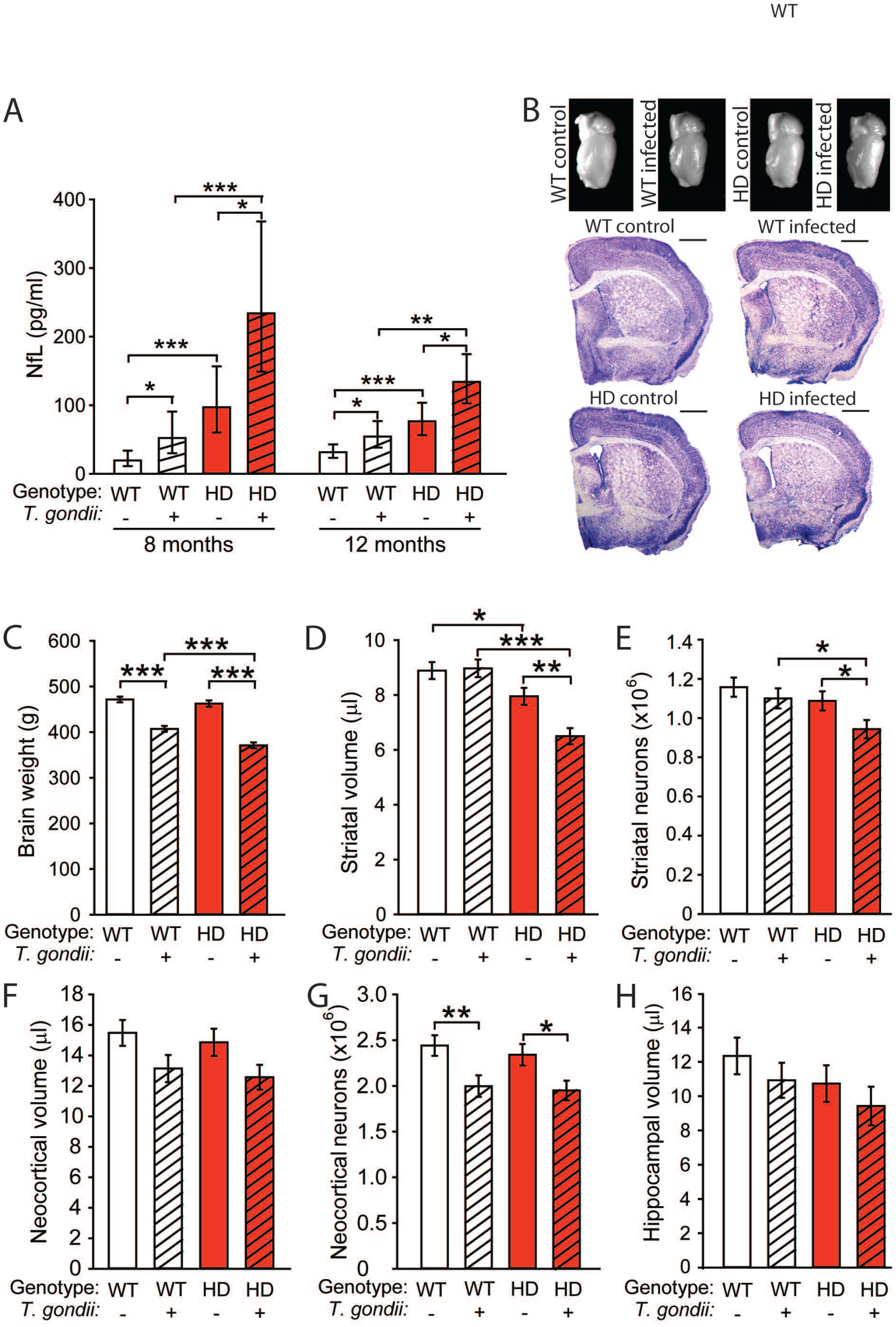
*T. gondii* exacerbates neurodegeneration in YAC128 mice. Mice were infected at 2-months of age. Blood was collected at indicated time points for NFL analyses, and neurodegeneration evaluated at 12-months of age. **A.** Plasma NFL is increased by *T. gondii* and HD at 8 and 12 months. Data are shown as the mean ± 95% CI; n = 6-9 and 10-15 at 8 and 12 months, respectively. **B.** Representative mouse brains (top) and thionin-stained sections (bottom). Scale bars = 1 mm. **C.** *T. gondii* infection decreases brain weights in wild-type (WT) and YAC128 mice. **D.** Non-infected HD mice demonstrate striatal atrophy compared with non-infected WT. *T. gondii* promotes striatal atrophy in HD, but not WT mice. **E.** *T. gondii*-infected HD mice have fewer striatal neurons compared to non-infected HD and infected WT. n = 6. **F.** Neocortical volume is decreased by *T. gondii* infection in WT and HD mice (main effect of infection p=0.0106). **G.** *T. gondii* infection decreases cerebral neocortical neuron estimates in WT and HD mice. **H.** HD or infection do not significantly impact hippocampal volumes. **B-H.** n=9-12. Data are shown as means ± SE. *p < 0.05, **p < 0.01, ***p < 0.001.

To determine if *T. gondii* infection affects HD-associated phenotypes, we measured motor performance and body weight. As expected non-infected HD mice demonstrated declines in performance as compared to non-infected WT mice. Further, *T. gondii* infected WT mice demonstrated declines in motor endurance from 9-months of age (**Fig. 2A**). YAC128 HD mice are reported to gain weight in adult life compared to WT controls [49]. Consistent with this non-infected HD mice gained weight compared to WT controls. Infection decreased body weight gains in HD mice, but had no effect on weight in WT mice (**Fig. 2B**).

**Figure 2.**
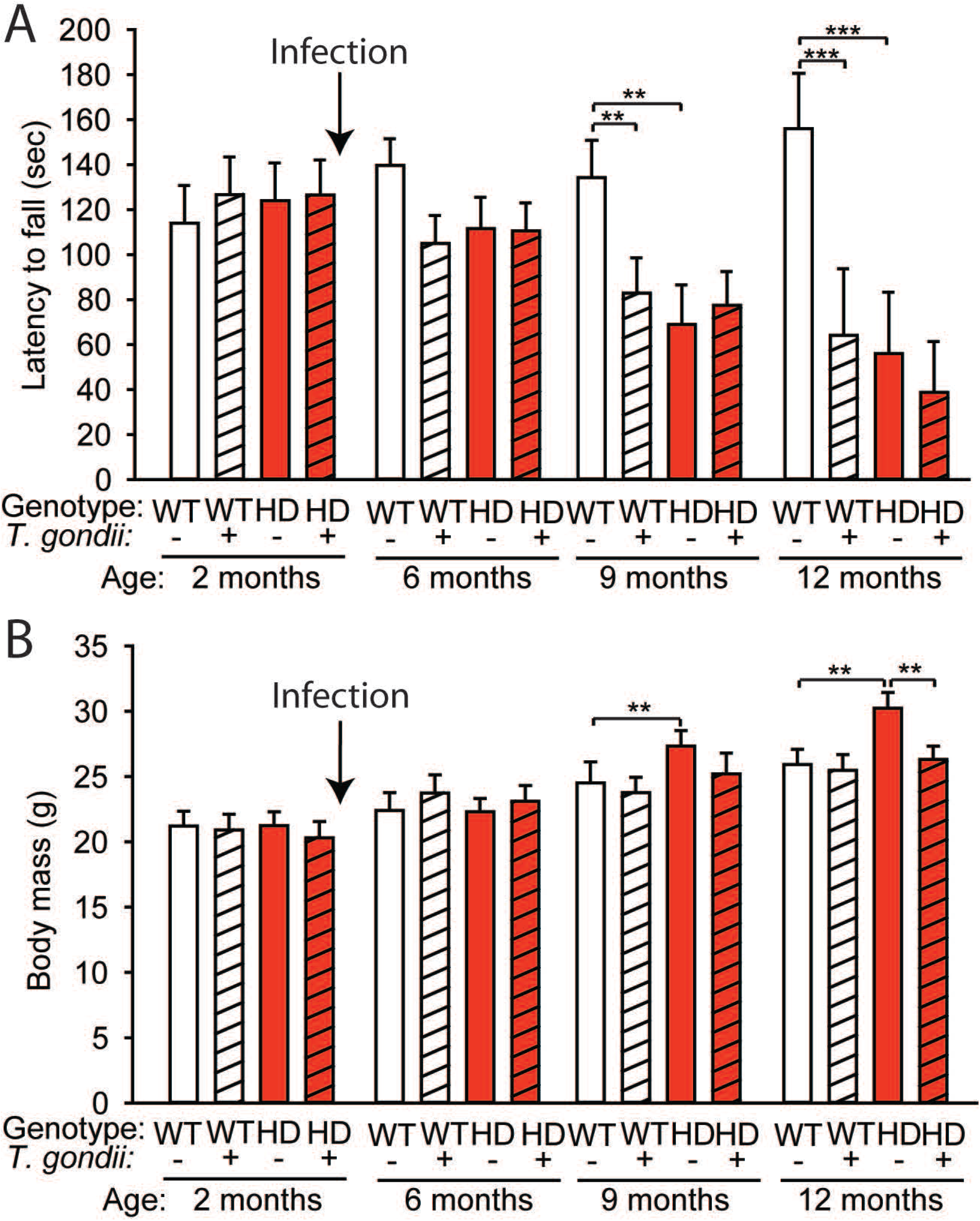
*T. gondii* induces progressive motor dysfunction and reduces body weight gain in YAC128 HD mice. Mice were infected at 2 months of age. **A.** Mouse motor endurance was tested using a rotating rod. *T. gondii* infection and HD lead to progressive declines in motor performance. **B.** *T. gondii* prevented weight gain in HD mice at 12-months of age. n=18-24. Data are shown as means ± SE. **p < 0.01, ***p < 0.001.

### YAC128 HD mice do not have a defect in parasite control or immune response

Differential infection-induced brain atrophy in WT and HD YAC128 mice could result from differences in parasite control. To address this we evaluated brain parasite cyst burdens (**Fig. 3**) and immune responses to infection (**Fig. 4**). At 8-months of age there were no differences in brain regional cyst burdens (**Fig. 3A**) or the number of parasite genomes (**Fig. 3B**) between WT and HD mice. *T. gondii* has two-life stages, a more virulent tachyzoite phase, which is characterized by rapid proliferation and expression of *SAG1*, and the bradyzoite phase, during which the parasites slowly replicate and express *BAG1*. Even though total parasite burden was not changed, we assessed whether HD mice had more tachyzoites, which would potentially induce more inflammation and injury. However, there were no differences in relative expression of life-stage markers between infected WT and HD mouse brain regions at 8-months of age (**Fig. 3C-D**). Further, at 12-months of age there were also no differences in cerebro-cortical brain parasite cyst numbers (**Fig. 3E**), or expression of *BAG1* (**Fig. 3F**) and *SAG1*(**Fig. 3G**). Therefore, greater parasite burdens in HD mice were not found.

**Figure 3.**
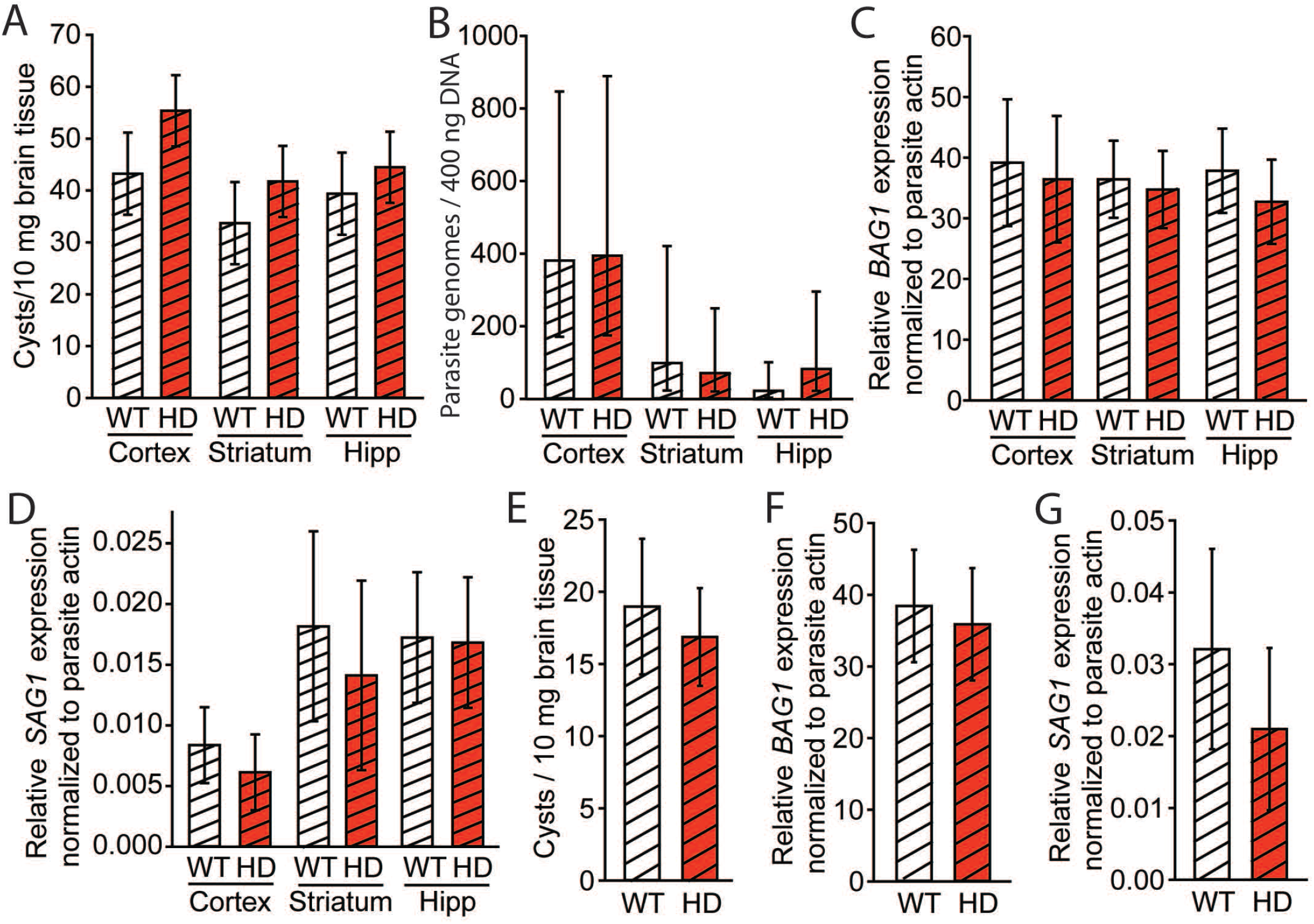
Equivalent *T. gondii* parasite load in WT and YAC128 mice. Mice were infected at 2 months of age and parasite loads determined at 8 (**A-D**) and 12 (**E-G**) months of age. **A.** Brain regional toxoplasma cyst counts are equivalent in infected WT and HD mice. n=6-9. **B.** Toxoplasma genome equivalents do not differ between infected WT and HD mice. n=6-9. Data are shown as means ± 95% confidence intervals. **C-D.** Bradyzoite-specific BAG1 (**C**) expression is equivalent in infected WT and HD mice. n = 7. **D.** Tachyzoite-specific SAG1 expression is equivalent in infected WT and HD mice. n = 7. **E.** Cerebral cortical toxoplasma cysts counts are the same valent at 12 months of age. n=13 WT and 18 HD mice. **F-G.** WT and HD mice demonstrate equivalent cerebro-cortical bradyzoite (**F**) and tachyzoite (**G**) parasite loads. n = 7. **A-D**, Hipp=hippocampus. Unless otherwise indicated data are shown as the means ± SE.

**Figure 4.**
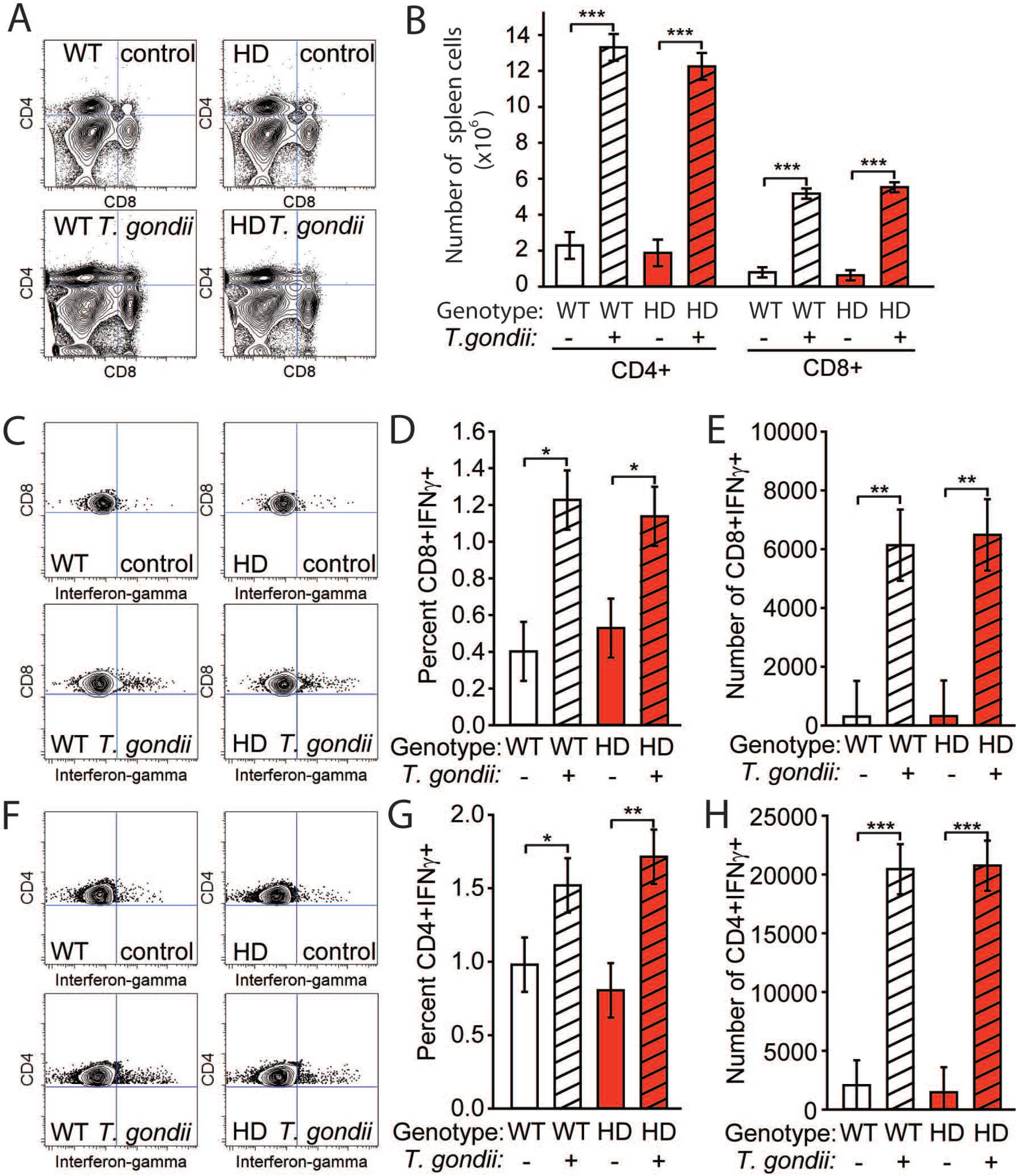
HD does not alter CD8+ and CD4+ T-lymphocyte responses to *T. gondii*. Mice were infected at 2 months; analyses were completed at 12-months of age. **A.** Spleen CD4+ and CD8+ T cells quantification. Representative flow cytometry contour plots of CD4+ (y axis) and CD8+ (x axis) populations. **B.** Total CD4+ and CD8+ spleen cells increases in infected mice. n = 6. **C.** Representative flow cytometry plots of IFNγ production by splenic CD8+ T cells. **D.** Percentages of IFNγ-producing CD8+ T cells are increased by infection in WT and HD mice. **E.** Total numbers of CD8+IFNγ+ T cells are increased by infection. **F.** Representative flow cytometry plots of IFNγ production by splenic CD4+ T cells. **G.** Percentages of IFNγ-producing CD4+ T cells are increased by infection in WT and HD mice. **H.** Total numbers of CD4+IFNγ+ T cells are increased by infection. **A, C and F:** top left = non-infected wild type, top right = non-infected HD, bottom left = infected wild type, bottom right = infected HD. n = 6. Data are shown as means ± SE. *p < 0.05, **p < 0.01, ***p < 0.001.

To evaluate immune *T. gondii* control we first determined the absolute number of splenic CD4+ and CD8+ T cells in 12-month-old mice. As expected T-cell numbers were increased by chronic-latent infection in WT mice. HD-infected mice showed the same response T-cell responses (**Fig. 4A-B**). IFNγ is critical for control of *T. gondii* infection [50]. We therefore measured T-cell IFNγ production in 12-month-old mice (**Fig. 4C-H**). Percentages (F_(1,20)_ = 9.66, p = 0.0055) and absolute numbers (F_(1,20)_ = 24.15, p < 0.0001) of splenic CD8+ IFNγ+ T-cells were increased infected mice (**Fig. 4C-E**). However, there were no differences in response between WT and HD mice. We also quantified splenic IFNγ-producing CD4+ T-cells as these are also important for long-term control of *T. gondii* [51, 52]. Similar to CD8+ cells, CD4+ T-cells had increased frequency of IFNγ production (F_(1,20)_ = 15.34, p = 0.0009) and increased total numbers of splenic CD4+ IFNγ+ cells (F_(1,20)_ = 78.15, p < 0.0001), but there were no differences between WT and HD mice (**Fig. 4F-H**). Therefore, differences in critical long-term immune responses to *T. gondii* infection were not found between WT and YAC128 HD mice.

### Altered peripheral kynurenine pathway metabolism in WT and YAC128 HD mice with *T. gondii* and HD in mouse brain

The kynurenine pathway of tryptophan degradation (KP) is implicated in HD progression [31, 32]. Furthermore, immune responses to *T. gondii* infection include activation of tryptophan metabolism as a means to limit this amino acids availability to the parasite, which it cannot synthesize. To investigate a potential role of the peripheral KP in latent *T. gondii*–induced neurodegeneration, we measured KP metabolites in blood at 2, 5, 8 and 12 month time points (**Fig. 5**). Tryptophan levels were not altered by infection or HD (**Fig. 5A**). However, there was a time-dependent increase in kynurenine with *T. gondii* infection (F_(3,114)_ = 7.85, p < 0.0001), and a HD-T. *gondii* interaction (F_(1,114)_ = 8.08, p = 0.0053) on kynurenine levels (**Fig. 5B**). The toxic metabolite 3-hydroxykynurenine was increased by *T. gondii* infection in WT and HD mice (F_(3,114)_ = 3.82, p = 0.012) at 8-months (**Fig. 5C**). However, neither infection nor HD resulted in changes in kynurenic acid levels (**Fig. 5D**). Therefore, infection resulted in different peripheral KP metabolism responses in WT and HD mice.

**Figure 5.**
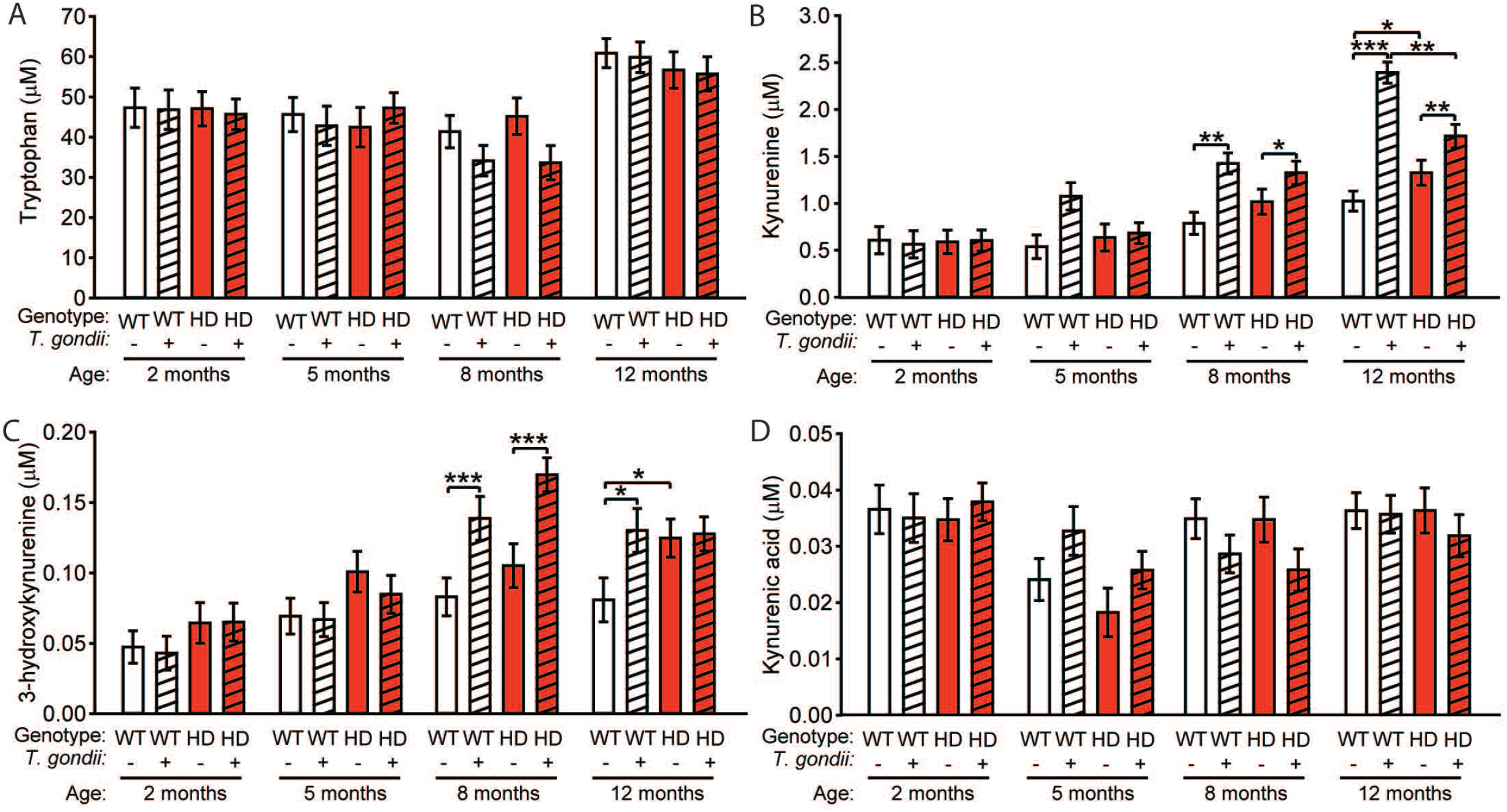
*T. gondii* and HD alter the peripheral kynurenine pathway in mice. Mice were infected at 2-months of age, and blood analyses completed at 2 (pre-infection), 5, 8 and 12 months of age. **A.** Tryptophan levels are not different between groups. **B.** Age-dependent increase in kynurenine in infected (WT and HD), and non-infected HD mice. **C.** *T*. gondii-infected WT and HD mice **D.** Kynurenic acid levels are not different between groups. n = 8-11. Data are shown as mean ± SE. *p < 0.05, **p < 0.01, ***p < 0.001.

### *T. gondii* increases soluble mHTT in YAC128 HD brain

Mutation of the huntingtin gene *(mHTT)* is the proximal cause of HD and results in expression of mHTT protein. Mutant huntingtin protein exists in equilibrium between soluble and aggregated forms [28]; however, soluble species are more toxic [53–55]. Therefore, we investigated the possibility that the parasite could promote HD by increasing *mHTT* gene expression or mHTT accumulation (**Fig. 6**). We quantified *mHTT* mRNA in three brain regions at 8 and 12 months of age in infected and non-infected HD mice and found no differences (**Fig. 6A-B**). We then measured soluble mHTT in cerebral cortex of HD mice at 12-months of age and found that soluble mHTT was significantly increased by infection (F_(1,13)_ = 7.32, p = 0.0180) (**Fig. 6C**).

**Figure 6.**
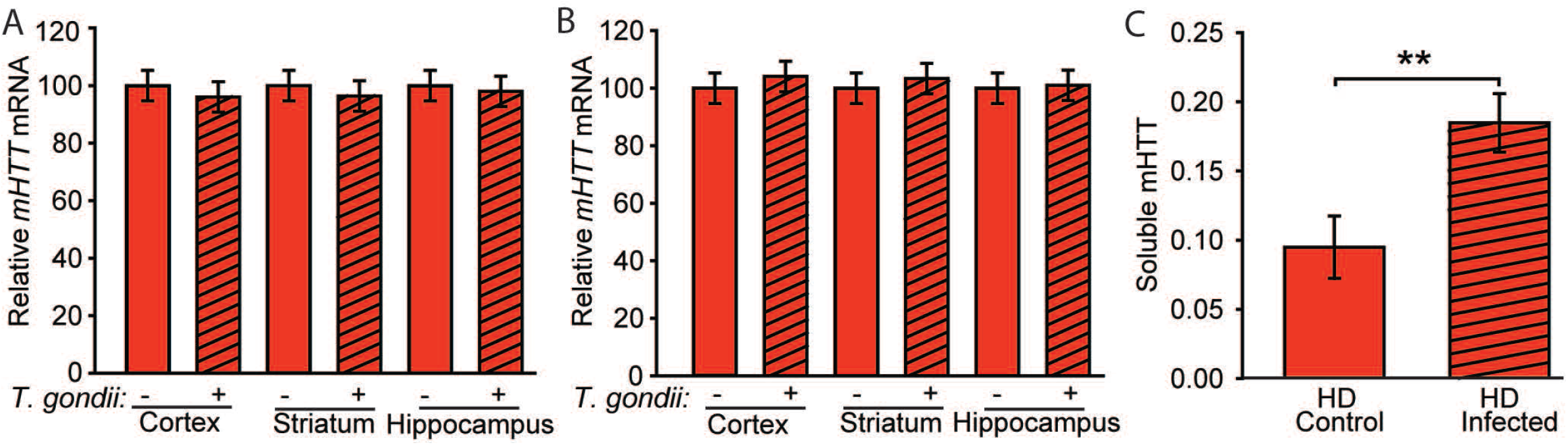
*T. gondii*infection increases soluble mHTT in YAC128 HD mice. Mice were infected at 2 months of age. **A-B.** *T. gondii* infection does not significantly alter *mHTT* at 8-months (**A**) or 12-months (**B**) of age. Transcript was quantified in cerebral cortex, striatum and hippocampus. n = 6. **C.** Soluble mHTT measured by the FRET ratio between two antibodies targeted to the N-terminal region of the protein. *T. gondii* significantly increased soluble mHTT at 12 months of age in the cerebral cortex. n = 7 non-infected HD and 8 infected HD mice. Data are shown as the mean ± 95% CI. **p < 0.01.

## Discussion

The age of onset of human HD is inversely proportional to the extent of CAG expansion, such that individuals with small expansions develop late-onset disease whereas large expansions result in childhood onset [56]. The extent of CAG expansion explains 50-70% of the variance in the age of HD onset, and the remaining variance is explained by environmental and non-HTT genetic factors. In a large Venezuelan study of HD patients, unknown environmental factors were estimated to contribute ~60% of the variance in age of onset after controlling for CAG mutation size [5]. Therefore, evidence suggests that environmental factors are significant modifiers of HD, although these modifiers are poorly understood. Chronic latent CNS infections, with agents such as *T. gondii* and HSV1, are putative modifiers of neurological disease. They promote low-grade inflammation that is needed to maintain control of the pathogen, and last for the post-infection life of the host. Mice are naturally infected with *T. gondii*, so they provide a suitable system for modeling the human infection. To study potential interactions between HD and *T. gondii*, we developed a model in which mice develop chronic latent infection to approximate the human infection, as occurs in immunocompetent individuals. We found that mice on the FVB/NJ background developed fully latent infection, enabling us to study the effects in YAC128 HD mice. We then demonstrated that latent parasite infection significantly increased neurodegeneration in HD mice and elevated toxic soluble mHTT. These changes could not be attributed to reduced parasite control or an altered immune response in HD mice. We did not observe decreased motor endurance in infected-HD mice. However, body weight differences confound motor endurance testing in HD mice [57] and the infected-HD group had significantly lower weights compared to non-infected HD mice (**Fig. 2**). Therefore, our data provide proof of principle that a latent brain infection can promote progression in a mouse model of HD.

*T. gondii* seropositivity is associated with increased risk of psychiatric disorders including schizophrenia and depression [58–60], but the nature of this relationship is debated [13, 18]. The current findings, and an earlier study [39], show that HD mice respond differently to infection with the moderately pathogenic ME49 strain of *T. gondii*, as compared with control mice. We found that latent infection also had adverse effects in control mice but that there is preferential susceptibility to the infection in HD mice. It is currently unclear how these findings relate to individuals with HD, who would be predicted to have toxoplasma infection prevalence rates that are similar to those of the general population. Because of differences in lifespan, humans may be infected for decades with *T. gondii*, providing more time for the effects of low-grade inflammation to manifest. However, although our mice were infected with only 10 tissue cysts, brain tissue burdens were notably higher than that which occurs during human latent toxoplasmosis. Therefore, direct extrapolation of our findings to human HD is not possible. Human studies are needed to determine whether latent toxoplasmosis is an environmental modifier of HD progression. Despite this caveat, our study provides proof of principle that chronic latent toxoplasmosis in HD mice potentiates established markers of HD progression. Interestingly, a recent study reported that early-life infection with *T. gondii* alters a wide array of human genes and pathways implicated in the pathogenesis of neurological disorders including HD, Alzheimer’s disease, and seizure disorders [61]. In humans, the prevalence of *T. gondii* increases with age [6, 62]. Therefore, a major question is how quickly *T. gondii* can induce persistent neurological changes and whether early infection is required for exacerbation of neurological diseases that manifest later in life.

Chronic *T. gondii* infection resulted in similar amounts of cortical atrophy in WT and HD mice. These effects are consistent with reports of motor impairment, memory deficits, and olfactory dysfunction resulting from human *T. gondii* infection [63–66]. They are also consistent with a report in humans that *T. gondii* reduces cortical grey matter volume [67]. Cortical atrophy in WT mice shows that *T. gondii* infection can result in neurodegeneration. Importantly, however, striatal volume and estimation of the number of striatal neurons demonstrated significant potentiation of HD neurodegeneration by infection, consistent with YAC128 HD mice being specifically vulnerable to the effects of infection, rather than showing an additive effect of infection and HD. This interpretation is supported by our important finding of elevated cortical mHTT in infected HD mice. In addition, increases in NFL, a marker of active neurodegeneration [47, 48], were greater in HD mice than in WT mice after infection. Given that differences in parasite burden were not present, our results suggest an increased vulnerability to the presence of *T. gondii* or to the associated inflammatory response in this mouse model of HD.

*T. gondii* infection amplified degeneration that is characteristic of HD, namely striatal atrophy. The vast majority (~95%) of striatal neurons are GABAergic medium spiny neurons. Recently, it has been reported that *T. gondii* alters presynaptic localization of GAD67, a marker for GABAergic neurons, which indicates that infection affects GABAergic synapses and thus could exacerbate HD-induced dysfunction in striatal medium spiny neurons [68]. In HD, glutamatergic cortico-striatal circuits are dysfunctional [69, 70]. During *T. gondii* infection, there is elevated extracellular glutamate caused by downregulation of the glutamate transporter, GLT-1 [71]. HD patients and mice also display reduced glutamate uptake and decreased GLT-1 expression [72–74]. These findings therefore suggest that one potential mechanism of HD potentiation by *T. gondii* involves promotion of dysfunction and glutamate and GAGAergic synapses.

Mutant HTT activates microglial cells and mHTT-expressing monocytes have an altered response to LPS treatment [27, 75]. HD mice also have increased microglial activation in response to peripheral LPS challenge [76]. One potential mechanism by which inflammation may alter HD is through activation of the kynurenine pathway (KP). HD patients have increased 3-hydroxykynurenine, which is produced as a result of kynurenine monooxygenase (KMO) oxidation of kynurenine, the second step in the KP [32, 77]. In our study, we also observed elevated kynurenine and 3-hydroxykynurenine in blood of *T. gondii*–infected mice earlier than these metabolites were elevated in HD mice without infection. Activation of the KP coincided with more pronounced neurodegeneration in HD mice. We propose a model whereby mHTT alters the initial responses to inflammatory insult resulting in an altered trajectory of the disease through KP activation and other mechanisms. Future studies will investigate the relationship between peripheral and brain KP metabolism.

Cytokines including IL-6 and TNFα, that are characteristic of neuroinflammation, reduce the clearance of misfolded proteins such as Aβ and tau [44, 78-80]. *T. gondii* infection, which also results in elevated IL-6 and TNFα, induces Aβ deposition and tau hyperphosphorylation that are associated with behavioral deficits in mice [25, 38]. Interestingly, one mechanism by which *T. gondii* evades clearance is by inhibiting autophagy [81, 82], and suppression of autophagy is sufficient to induce neurodegeneration [83, 84]. Therefore, as these cytokines are also elevated in HD, environmental factors that increase neuroinflammation such as *T. gondii*could impair clearance of mHTT. Macroautophagy is critical for clearing mHTT, but these pathways are also dysfunctional in HD [63, 85, 86]. Promoting autophagy is protective in HD models, and polymorphisms in autophagy-related genes are associated with earlier age of onset in HD [85, 87, 88]. We found that *mHTT* mRNA expression was unchanged by *T. gondii*.However, an important finding of our work is that there was increased soluble mHTT in infected HD mice. Therefore neuroinflammation, which alters the clearance of misfolded proteins, may represent a key mechanism by which latent infection exacerbates HD progression by inducing accumulation of toxic soluble mHTT. Of note, kynurenine is reported to activate the mammalian target of rapamycin (mTOR) in T cells, which decreases autophagy [89]. Therefore, increased mHTT accumulation in our study is consistent with elevated KP activation resulting from both latent *T. gondii* infection and HD. Based on our studies, more work is needed to understand how inflammation and/or KP activation resulting from latent brain infections affects mHTT expression, aggregation, and clearance.

In conclusion, we established a novel system for studying the interaction between latent toxoplasmosis and HD, a monogenetic protein-misfolding disease. Our findings corroborate other studies showing that latent *T. gondii* infection has adverse effects on the aging brain in WT mice. We additionally demonstrated that YAC128 HD mice demonstrate specific vulnerability to this infection with promotion of key disease markers. This study provides a framework for understanding how neuroinvasive infections may promote neurodegeneration in HD and related neurodegenerative diseases, as well as brain aging.

## References Supporting Information

**Table S1.** Mouse deaths in reported studies. Listed are all mouse deaths or exclusions across all studies used to generate the data in this manuscript. Exclusion occurred only in response to mice that were fighting. CON = vehicle treated, INF =*T. gondii* infected.

| Sex | Genotype | Treatment | Age (months) | Sex | Genotype | Treatment | Age (months) |
| --- | --- | --- | --- | --- | --- | --- | --- |
| Female | HD | INF | 3 | Female | WT | INF | 6 |
| Male | WT | INF | 3 | Male | WT | CON | 7 |
| Male | HD | CON | 3 | Male | WT | INF | 7 |
| Male | WT | INF | 4 | Male | HD | CON | 7 |
| Female | WT | INF | 4 | Male | WT | CON | 7 |
| Male | WT | INF | 4 | Female | HD | CON | 7 |
| Male | HD | INF | 5 | Male | WT | INF | 8 |
| Female | WT | INF | 5 | Female | HD | CON | 8 |
| Female | HD | INF | 5 | Male | HD | CON | 8 |
| Male | WT | INF | 5 | Male | WT | CON | 8 |
| Female | HD | INF | 5 | Female | WT | CON | 10 |
| Female | WT | CON | 6 | Male | WT | CON | 11 |
| Male | WT | CON | 6 | Female | HD | INF | 11 |
| Male | HD | CON | 6 | Female | WT | INF | 11 |
| Male | HD | INF | 6 | Female | HD | CON | 11 |
| Male | HD | INF | 6 | Female | HD | CON | 11 |
| Male | HD | CON | 6 | Male | HD | CON | 11 |
| Male | HD | CON | 6 | Male | WT | CON | 11 |
| Female | WT | INF | 6 | Female | WT | CON | 11 |
| Female | WT | INF | 6 | Male | HD | INF | 12 |

**Figure S1.**
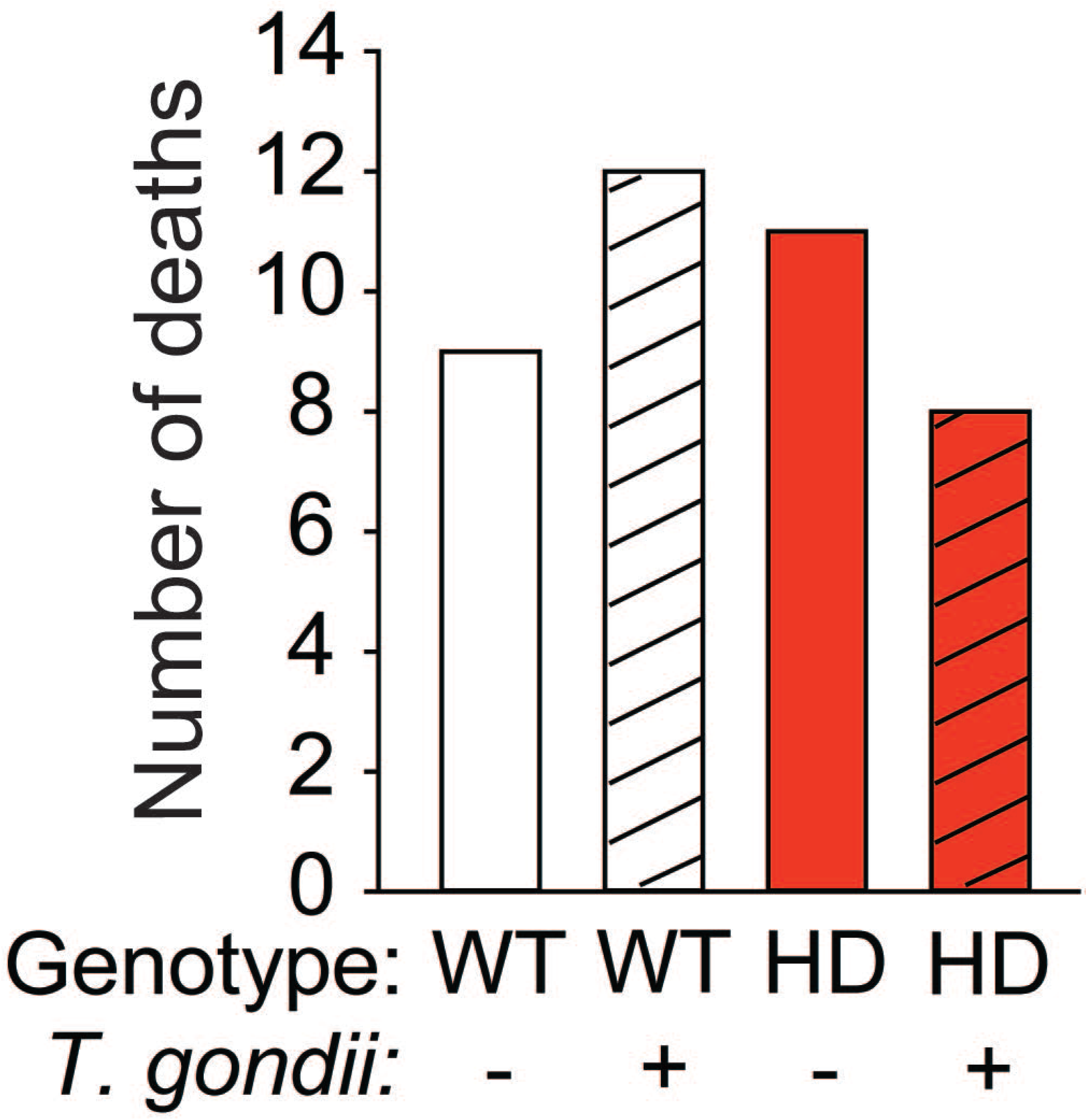
No differences in mouse mortalities among groups. Histogram shows the total number of spontaneous deaths that occurred across studies. There were no significant differences among the groups with respect to survival times.

